# Understanding C_4_ photosynthesis in Setaria by a proteomic and kinetic approach

**DOI:** 10.1101/2021.03.16.435684

**Authors:** Paula Calace, Tomás Tonetti, Ezequiel Margarit, Carlos M. Figueroa, Carlos Lobertti, Carlos S. Andreo, Mariel C. Gerrard Wheeler, Mariana Saigo

## Abstract

Plants performing C_4_ photosynthesis have a higher productivity per crop area related to an optimized use of water and nutrients. This is achieved through a series of anatomical and biochemical features that allow the concentration of CO_2_ around RuBisCO. In C_4_ plants the photosynthetic reactions are distributed between two cell types, they initially fix the carbon to C_4_ acids within the mesophyll cells (M) and then transport these compounds to the bundle sheath cells (BS), where they are decarboxylated so that the resulting CO_2_ is incorporated into the Calvin cycle (CC).

This work is focused on the comparative analysis of the proteins present in M and BS of *Setaria viridis*, a C_4_ model close relative of several major feed, fuel, and bioenergy grasses. The integration of kinetic and proteomic approaches agrees that the C_4_ compound malate is mainly decarboxylated in the chloroplasts of BS cells by NADP-malic enzyme (NADP-ME). Besides, NAD-malic enzyme (NAD-ME) located in the mitochondria could also contribute to the C_4_ carbon shuttle. We presented evidence of metabolic strategies that involve chloroplastic, mitochondrial and peroxisomal proteins to avoid the leakage of C_4_ intermediates in order to sustain an efficient photosynthetic performance.

**Highlight:** Proteomic and kinetic analyses show metabolic strategies involving chloroplastic, mitochondrial and peroxisomal proteins to maintain the C_4_ cycle performance in parallel to other metabolic pathways.

## Introduction

Climate change and future population food and fiber demands require the improvement of crop yields. A deeper understanding of the dynamic response of plants to changing environmental conditions is a key challenge to develop new crop varieties (Bayley-Serres *et al*., 2019). Furthermore, to enable future production systems to operate more sustainably, it is necessary to increase nitrogen and water-use efficiency (Mueller *et al*., 2012). The plants that perform C_4_ photosynthesis have a higher productivity per crop area, related to an optimized use of water and nutrients, due to the operation of a carbon concentration mechanism (CCM) (Sage, 2004). RuBisCO is a bifunctional enzyme that catalyses not only CO_2_ assimilation by carboxylation of ribulose-1,5-bisphosphate (RuBP), but also catalyses its oxygenation, generating 2 molecules of 3-phosphoglycerate (3PG) or one of 3PG and 2-phosphoglycolate (PG), respectively. The 3PG pool is used to regenerate RuBP and produce triose phosphate that will fuel carbohydrate biosynthesis. On the other hand, PG to be recycled must first be converted to pyruvate (Pyr) by a series of reactions collectively known as photorespiration. This metabolism is energetically costly and a quarter of the carbons are lost as CO_2_. PG production is higher with increasing leaf temperature, a condition that occurs in hot climates, with high irradiation and low evapotranspiration due to stomatal closure. This situation is aggravated by a photoinhibition condition, as the NADPH generated in the light reactions accumulates due to the drop in the reductive assimilation of CO_2_ in the Calvin cycle. Consequently, photosynthetic capacity is reduced by up to 30% in hot and arid environments (Schulze and Hall, 1982; Jordan and Ogren, 1984; Bauwe *et al*., 2010).

The increase of the carboxylation over the oxygenation reaction in the active site of RuBisCO has a penalty of lower rates of product release and hence diminished rates of carboxylation (Shih *et al*., 2016). Then, engineering efforts would be more profitable if oriented to develop CCMs that increased the local concentration of CO_2_ surrounding RuBisCO (Shih *et al*., 2016). Plants with C_4_ metabolism use, in addition to the enzymes commonly found in C_3_ plants, a set of enzymatic activities compartmentalised in two cell types that allow them to enrich the RuBisCO environment in CO_2_, thus reducing its oxygenase activity (Figure 1). First, primary CO_2_ fixation occurs in the outer mesophyll (M) compartment, where carbonic anhydrase (CA) converts CO_2_ into HCO_3_ ^−^, which is used by phosphoenolpyruvate (PEP) carboxylase (PEPC) to carboxylate one molecule of PEP (C_3_), thus producing oxaloacetate (OAA, C_4_). Unlike RuBisCO, PEPC has no affinity for O_2_. Depending on the species, the OAA is further transformed into malate (Mal, C_4_) or aspartate (Asp, C_4_), which are transported to the inner layer of cells known as bundle sheath (BS). These cells possess several characteristics of the Kranz anatomy that avoid their contact with ambient air: cell walls with reduced gas permeability and positions adjacent to the vascular bundle. Depending on the prevalent decarboxylating activity in green leaves, C_4_ species have been traditionally classified as NADP-malic enzyme (NADP-ME), NAD-malic enzyme (NAD-ME) or phosphoenolpyruvate carboxylase (PEPCK) subtypes. In NADP-ME subtype species, Mal is the C_4_ acid transported from M to BS and it is decarboxylated in plastids by NADP-ME. In NAD-ME and PEPCK species, Asp is transported between cells and further transformed back to Mal or OAA, which are decarboxylated by NAD-ME or PEPCK in mitochondria or the cytosol, respectively. The generated C_3_ acids return to the M, where they are used to regenerate PEP through the action of the enzyme pyruvate phosphate dikinase (PPDK, Figure 1). The decarboxylase selection affects many aspects of C_4_ photosynthesis beyond the biochemical pathway, such as BS and M ultrastructure, M to BS transport processes, leaf energetics, and photosynthetic efficiency (Hatch, 1987; Drincovich *et al*., 2011; Ghannoum *et al*., 2011). A growing body of evidence suggests that C_4_ photosynthesis can involve more than one decarboxylase in the same species, which would enable a high photosynthetic performance even in changing environmental conditions. It is currently proposed to subclassify C_4_ plants only as NADP- or NAD-ME subtype plants, as there is a clear line of demarcation based on the type of major decarboxylase, with variable contributions from PEPCK (Furbank, 2011; Wang *et al*., 2014). Thus, PEPCK could be considered as a complementary activity to Mal-decarboxylating enzymes, not as an independent C_4_ decarboxylation activity, which would provide additional mechanisms to balance energy between BS and M under different light conditions and to help lower metabolite gradients required for the C_4_ cycle functioning.

**Figure 1:**
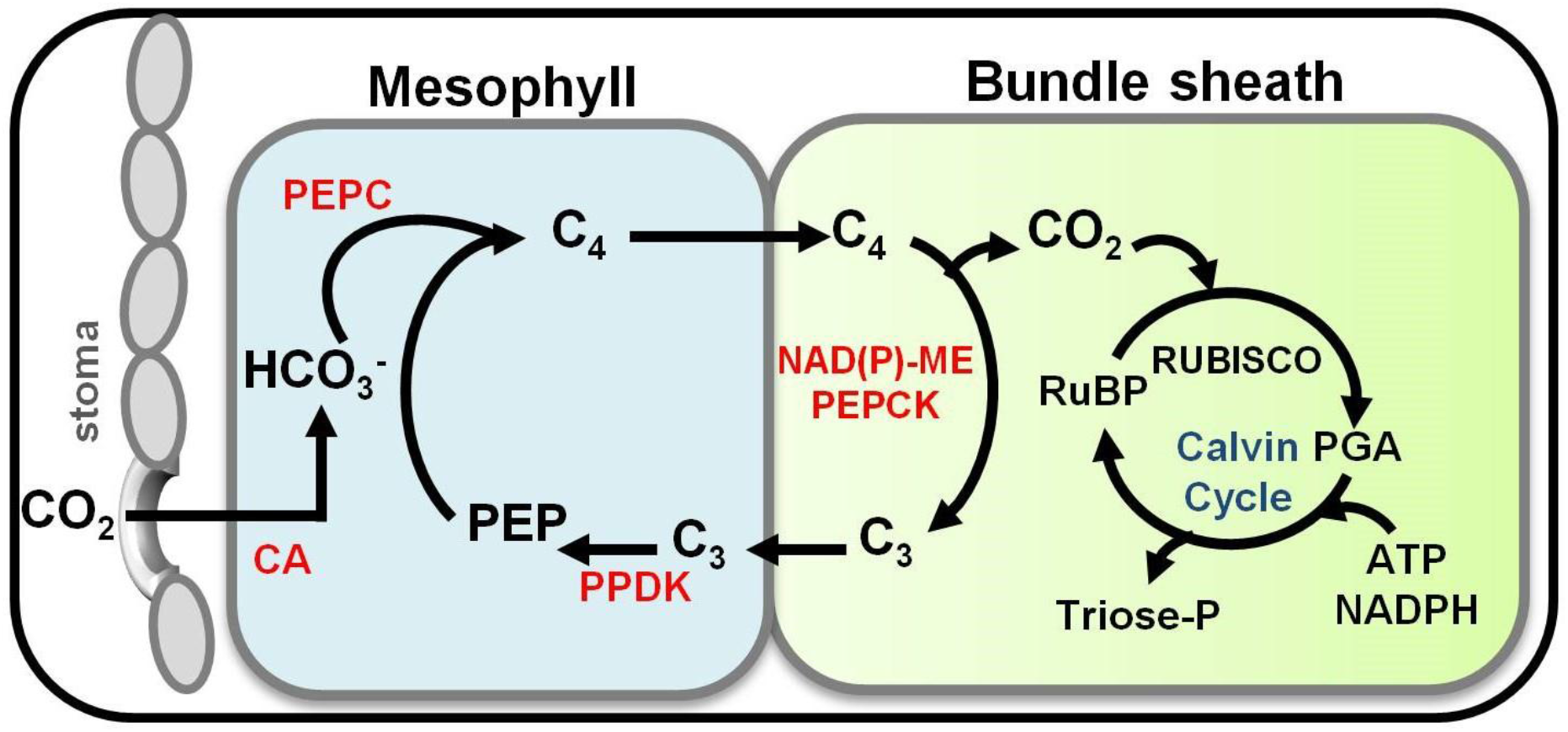
Schematic of the C_4_ photosynthesis pathway. In C_4_ plants the photosynthetic reactions are distributed between two cell types, they initially fix the carbon to C_4_ acids within the mesophyll cells and then transport these compounds to the bundle sheath cells, where they are decarboxylated so that the resulting CO_2_ is incorporated into the Calvin cycle. This cycle produces the enrichment of the RuBisCO environment in CO_2_, thus reducing its oxygenase activity. Names in black correspond to metabolites; CO_2_: carbon dioxide; HCO ^−^: bicarbonate; C: 4-carbon acid; C_3_: 3-carbon acid; PEP: phosphoenolpyruvate; RuBP: ribulose-1,5-bisphosphate; PGA: 3-phosphoglyceric acid. Names in red correspond to C_4_ genes; CA: carbonic anhydrase; PEPC: phosphoenolpyruvate carboxylase; PPDK: pyruvate/orthophosphate dikinase; NAD(P)-ME: NAD-dependent malic or NADP-dependent enzyme; PEPCK: phosphoenolpyruvate carboxykinase.

The enzymes involved in the C_4_ cycle are part of protein families, where the other members perform various housekeeping functions, and are therefore referred to as “C_3_-type”. The C_3_-type genes would have been the genetic basis for the emergence of the C_4_-type variants through duplication and neofunctionalization, which allowed C_4_ plant species to gain new enzymes without losing the previous functions. High expression levels, response to light, adequate compartmentalisation and optimal structural, catalytic and regulatory properties were the foundations for their new functionality (Sage, 2004; Edwards and Smith, 2010; Maier *et al*, 2011; Saigo *et al*, 2013a, Alvarez *et al*, 2019).

Among grasses (Poaceae), the subfamily Panicoideae includes very important species from an agronomic point of view, as they are used as raw material in the food and biofuel industry. Maize, sorghum and sugarcane all belong to this group and perform a NADP-ME type C_4_ photosynthesis. The subclade Paniceae is of particular interest as it includes species of NADP- and NAD-ME subtypes, such as *Setaria viridis* (green millet, C_4_-NADP-ME), *Setaria italica* (foxtail millet, C_4_-NADP-ME) and *Panicum virgatum* (switchgrass, C_4_-NAD-ME). *S. viridis* is a small plant, its life cycle lasts between 6 and 9 weeks, it has a small (510 Mb) and sequenced genome (Bennetzen *et al*., 2012), and robust transformation protocols have been developed for this species (Van Eck, 2018). These technical characteristics and its phylogenetic closeness to species of great agronomic importance have positioned *S. viridis* as a model within the C_4_ grasses (Acharya *et al*., 2017; Doust *et al*., 2019). *S. italica* is a forage crop and a very close relative of *S. viridis*, since it was developed in China from its domestication.

In this work, we analyse the C_4_ pathway of *S. viridis* underlining the metabolic context that supports the photosynthetic process. First, we show that BS enrichment, kinetic performance and regulation of NADP- and NAD-ME agree with a major role of chloroplastic NADP-ME in C_4_ cycle and a potential auxiliary role of mitochondrial NAD-ME. Second, we describe the differential proteomics of M and BS of chloroplasts and mitochondria, since they would have different contributions to the C_4_ carbon shuttle, depending on the site of Mal decarboxylation. Finally, we report the peroxisomal metabolic activities in M and propose novel roles in the context of the C_4_ photosynthesis. The comparison with maize emphasizes the need to characterize the metabolic strategies operating for C_4_ cycle in a variety of plant species in order to discriminate the core conserved characteristics from those species-specific. This knowledge help to understand more deeply the C_4_ photosynthesis, which is essential to face challenges on plant productivity improvement.

## Materials and methods

### Plant growth and harvest conditions

Seeds of *Setaria viridis* A10.1 were germinated on plates and then sown on soil. First, seed coats were removed with sandpaper to break the dormancy and to guarantee a high percentage of germination (Van Eck and Swartwood, 2015). Second, seeds were surface-sterilized with a mixture of 1% (w/v) sodium hypochlorite and 0.1% Tween-20, and then thoroughly washed with sterile water. Finally, sterile seeds were sown on 0.5X MS-agar plates and incubated at 28°C with irradiance of 120 μmol m^−2^ s^−1^ (16h light/8h dark) for 5 days. Seedlings were transferred to individual 8-cm pots containing Klasmann TSI substrate and irrigated with 1X Hoagland solution. Plants were grown in a Conviron Adaptis A1000 chamber with irradiance of 350 μmol m^−2^ s^−1^ (16h light/8h dark), at 28/22°C (day/night) and 50% relative humidity for 2 weeks. At this stage, plants usually had 8 fully developed leaves and inflorescences were not visible. Leaves 5 and 6 (counting from the bottom of the plant) were harvested at the middle of the photoperiod and immediately used to isolate mesophyll cells (M) and bundle sheath strands (BS).

### Separation of M and BS

M cells were isolated using the leaf rolling protocol (Covshoff *et al*., 2013) with the following modifications. The centre of each leaf was cut in two 5-cm segments and the midrib was removed to generate four segments per leaf. Samples were placed on an ice-cold glass and a plastic rod was rolled twice over the surface of each leaf segment to release the M content, which was rapidly collected using a pipette and dispensed into a 1.5 ml tube on liquid nitrogen.

BS strands were isolated using a method modified from John *et al*. (2014). As mentioned above, leaves were divided into four segments, which were further cut into 2-mm segments. The fragments were placed in isolation buffer (0.33 M sorbitol, 0.3 M NaCl, 0.01 M NaCl, 0.01 M EGTA, 0.01 M dithiothreitol, 0.2 M Tris-HCl pH 9.0, and 5 mM diethylthiophosphoryl chloride), and then pulsed for 10 s three times in a hand blender on low speed. The suspension was then filtered through a 60-μm mesh, and blending buffer (0.35 M sorbitol, 5 mM EDTA, 0.05 M Tris-HCl pH 8.0, and 0.1% (v/v) 2-mercaptoethanol) was used to return the BS material back into the blender. Homogenization at maximum speed for 1 min followed by filtering was repeated three times. Before the last filtration step, the suspension was strained by a homemade, coarser-pore strainer to better clean the final sample. Purified BS strands were placed on a paper towel stack to remove excess moisture and then frozen in liquid nitrogen.

### Total protein extraction

For protein extract preparation, 100 mg of Setaria M and BS samples were pulverized in the presence of 1 ml of extraction buffer, which contains 500 mM Tris-HCl pH 8.0, 50 mM EDTA, 700 mM sucrose, 100 mM KCl, 5% (w/v) poly-vinylpolypyrrolidone, 2% (v/v) 2-mercaptoethanol, and 1% (v/v) complete protease inhibitor cocktail (Roche). Then, 1 ml of phenol equilibrated to pH 8.0 was added and the mixture was incubated on ice and vortexed for 15 min before centrifugation at 10,000 x g for 10 min at 4°C. The organic fraction was collected and five volumes of 100 mM ammonium acetate in methanol were subsequently added. Proteins were left to precipitate for 24 h at −20°C. Samples were centrifuged at 10,000 x g for 20 min at 4°C, and supernatants were removed. Resulting pellets were washed twice with 100 mM ammonium acetate/methanol. Finally, each pellet was suspended in 250 μl of sample buffer (12 mM Tris-HCl pH 6.8, 0.4% (w/v) SDS, 1.5% (w/v) dithiothreitol and 5% (v/v) glycerol) and boiled twice for 5 min. Protein quantification was performed by the Bradford method (Bradford, 1976) using bovine serum albumin as standard.

Protein extracts were checked by Western blot assay in order to verify a correct and efficient separation of cells. For this, M and BS protein samples were run on 10% (w/v) polyacrylamide gels for SDS-PAGE (Laemmli, 1970) and then electroblotted onto nitrocellulose membranes for inmunoblotting according to Burnette (1981). Anti- RuBisCO and PEPC antibodies were used for detection. Bound-antibodies were located by linking to alkaline phosphatase-conjugated goat anti-rabbit IgG according to manufacturer’s instructions (Sigma).

Finally, proteins (35 μg per well) were loaded on a 10% (w/v) polyacrylamide gel for SDS-PAGE (Laemmli, 1970) and then run for 1 cm. Bands were visualized by staining with colloidal Coomassie Brilliant Blue G-250 and subsequent incubation with a 30% (v/v) methanol solution, and then cut with a sterile scalpel (Leonardi *et al*., 2015). Four pools of each sample (M and BS) were processed as biological quadruplicates.

### Mass spectrometry analysis of protein samples

Samples were digested with trypsin (Promega) and cleaned with ZipTips C18 (Milipore). The resulting peptides were analyzed by nanoHPLC coupled to a mass spectrometer with Orbitrap technology. A Thermo Scientific EASY-nLC 1000 chromatograph was used to separate protein complexes with a high degree of resolution using a reversed-phase column (Easy-Spray PepMap RSLC C18 column-3 μm, 100 A, 75 μm × 150 mm; Thermo Scientific) at 35°C. The injection volume was 4 μl. Aqueous and organic phases were 0.1% (v/v) formic acid in water or in acetonitrile, respectively. A two-step gradient of 5-35% (for 100 min) and 35-100% (for another 5 min) linear increment of the organic phase was used. The flow rate was kept at 200 nl min^−1^ in all steps. An ionizer with a spray voltage of 2.75kV was used to electrospray the eluted peptides. The configuration of the equipment allows peptide identification to be carried out at the same time as the peptides are separated by chromatography, obtaining Full MS (resolution: 70,000 FWHM) and MS/MS (resolution: 17,500 FWHM). A method that performs the highest number of measurement cycles per unit time was used. In each cycle the equipment performs a Full MS and then MS/MS to the 15 peaks with the best noise signal in that cycle, with a dynamic exclusion range to prevent the same peak from being fragmented more than once in the same elution peak of the chromatogram.

The identification and abundance estimation of each protein was performed with the Proteome Discoverer 2.2 program (Thermo Scientific) run against the *Setaria viridis* UP000298652 (UniProt) database. Peptides with a high level of confidence were considered. Search parameters were adjusted for an error tolerance equal to 0.05 Da for the fragment ions and 10 ppm for the parent ions. Oxidation of methionine and carbamidomethylation of cysteine residues were selected as dynamic and static modifications, respectively. Data was analyzed and plotted using Perseus 1.6.6.0 software (Tyanova *et al*., 2016). Missing values were replaced by the minimum value of the set of protein abundances in quadruplicate samples with a single value or none determined. Statistical analysis (Student’s t-test) was performed to compare the abundances of different proteins identified in each sample (M and BS).

### Bioinformatic analysis

To characterize the proteins of interest, Uniprot identifiers were assigned to those of Phytozome using the *Setaria viridis* v2.1 database (https://phytozome.jgi.doe.gov/pz/portal.html#!info?alias=Org_Sviridis_er). A comprehensive search for genetic information including orthologs assigned to *Arabidopsis thaliana*, *Zea mays*, and *Oryza sativa* was performed.

Functional categories were determined using MapMan X4 and Mercaptor 4.2 tools (Schwacke *et al*., 2019). For this, UniProt identifiers were converted to Ensembl plant identifiers using BioMart 0.7 (Kinsella *et al*., 2011; Howe *et al*., 2020).

The putative subcellular location of each protein was inferred by the identification of its ortholog in proteomic databases of maize chloroplasts (Friso *et al*., 2010), Arabidopsis mitochondria (Fuchs *et al*., 2020) and Arabidopsis peroxisomes (Pan and Hu, 2018). The analysis of the primary sequence by TargetP (Almagro Armenteros *et al*., 2019), further confirmed the chloroplastic or mitochondrial location of a number of proteins and expanded the assignment to others (Supplementary Table 1).

### Cloning of NADP-ME and NAD-ME isoforms

cDNAs encoding C_4_-NADP-ME (Seita.5G134300), NAD-ME1 (Seita.2G322000), and NAD2-ME2 (Seita.9G200600) were amplified by RT-PCR using RNA extracted from *Setaria italica* leaves with Quick-Zol (Kalium Technologies). The concentration and integrity of the preparations were assayed by 2% (w/v) agarose gel electrophoresis. One μg of total RNA was reverse transcribed using M-MLV (Promega) and oligodT as primer. Then, amplification was made using Phusion High-Fidelity DNA Polymerase(Thermo Fisher Scientific) and specific primers. In the case of C_4_-NADP-ME, the oligonucleotide pair: NdeIC4NADP-for (5′-GTGCAG<u>CATATG</u>GCGGTAGGC-3′) and SalIC4NADP-rev (5′-AGCGGTGACAAC<u>GTCGAC</u>CAAAAC-3′) was used. NAD-ME1 was amplified using NheINAD1-for (5′-TGC<u>GCTAGC</u>CCCGTCGTCC-3′) and SacINAD1-rev (5′-GCAAACA<u>GAGCTC</u>TCTAGTCTGTC-3′), while NAD2-ME was amplified using NheINAD2-for (5′-G<u>GCTAGC</u>TGCATCGTGCAC-3′) and XhoINAD2-rev (5′-AGA<u>CTCGAG</u>ATTTATTTGTCGCTC-3′). The oligonucleotides were designed to introduce restriction sites (underlined) to facilitate subcloning into the expression vector. PCR products were ligated into the pGEMT-easy vector (Promega) and then subcloned into the expression vector pET-28a, which yields recombinant proteins fused to hexahistidine tail (Novagen). The correct cloning was confirmed by sequencing.

### Expression and purification of recombinant enzymes

The recombinant plasmids containing the inserts of C_4_-NADP-ME, NAD-ME1 and NAD-ME2 were used to transform *Escherichia coli* BL21 (DE3). Bacteria were grown in autoinduction medium inducing expression (Studier, 2005). Fusion proteins were purified using Ni^2+^-containing His-Bind columns (Novagen). Purified proteins were desalted and concentrated by ultracentrifugation (Amicon) using buffer TMG (50 mM Tris-HCl pH 8.0, 10 mM MgCl_2_, and 10% (v/v) glycerol). The purity and integrity of the recombinant proteins were analyzed by SDS-PAGE revealed with Coomassie dye. Protein concentration was determined using the Bradford method (Bradford, 1976). The purified enzymes were immediately stored in small aliquots at −80°C in buffer TMG.

### NADP-ME and NAD-ME activity assays

NADP-ME activity was determined at 30°C in a Jasco V-730 spectrophotometer using a standard reaction mixture containing 50 mM Tris-HCl pH 8.0; 10 mM MgCl_2_; 0.5 mM NADP^+^; and 10 mM Mal. NAD-ME activity was measured using a reaction mixture containing 50 mM MES-NaOH pH 6.5; 10 mM MnCl_2_; 0.5 mM NAD^+^; and 10 mM Mal. The optimal pH for the reactions was determined using different buffers as follows: 50 mM sodium acetate-acetic acid (pH 4.5−5.5), 50 mM MES-NaOH (pH 6.0−6.5), 50 mM MOPS-KOH (pH 7.0) and 50 mM Tris-HCl (pH 7.5−8.0−8.9).

The kinetic characterization of the selected enzymes was performed by varying the concentration of one of the substrates (Mal or NADP^+^/NAD^+^), while keeping the level of the other substrate at a fixed and saturating concentration. All kinetic parameters were calculated by fitting to the Hill equation (Detarsio *et al*., 2003; Saigo *et al*., 2013b) in at least triplicate determinations. Since the true substrates of ME are the free forms not complexed with metal ions, the data were analyzed considering the free concentrations of Mal and NADP^+^ or NAD^+^ in the test medium. The following dissociation constants (*K*d) values for the metal-substrate or metal-cofactor complexes were used: Mg^2+^-Mal, 28.2 mM; Mn^2+^-Mal, 20.0 mM; Mg^2+^-NADP^+^, 19.1 mM; and Mn^2+^-NAD^+^, 12.9 mM (Grover *et al*., 1981).

By assaying different compounds as potential inhibitors or activators of enzyme activity, NADP-ME (or NAD-ME) activity was measured in the presence of 0.5 or 2 mM of each effector (citrate, fumarate, succinate, OAA, fructose 1,6-bisphophate, PEP, α-ketoglutarate, Glu, Ala, and Asp) and non-saturating concentrations of Mal, NADP^+^ or NAD^+^. In the case of CoA and acetyl-CoA, measurements were performed at a 20 μM concentration of each effector. Inhibition assay of NADP-ME by high Mal concentration was carried out at pH 7.0 (Detarsio *et al*., 2007).

The reductive carboxylation of Pyr (reverse reaction) was tested in various buffer systems (pH 6.5−8.0) containing different concentrations of Pyr (0.1−50 mM), NADPH or NADH (0.1−0.2 mM), NaHCO_3_ (15−30 mM) and metal cofactor (10 mM MnCl_2_ or MgCl_2_).

## Results and Discussion

### General overview of M and BS proteomes

In order to analyse the proteins enriched in M and BS of Setaria leaves, we separated M cells by the leaf rolling method (Covshoff *et al*., 2013) and BS strands by a method of blending and filtering (John *et al*., 2014). M and BS proteins obtained by a phenol-based extraction (Leonardi *et al*. 2015) show clearly different patterns in Coomassie-stained gels (Supplementary Figure 1A). To further confirm the quality of the separations, we checked the enrichment of PEPC in M and RuBisCO in BS by Western blots (Supplementary Figures 1B and 1C).

According to the peptide identification and quantification by mass spectrometry, 1,385 proteins were identified in the quadruplicates of M and BS samples. Statistical analysis shows that 945 proteins (68% of total detected) are differentially expressed in M and BS cells in a significant way (FDR-*p*-value < 0.05), while 581 and 364 proteins are more abundant in M and BS, respectively (Supplementary Table 1, Figure 2A). Furthermore, 71 and 81% of proteins enriched in BS and M increase their abundance to more than double (log_2_(BS/M)>1 and <−1, respectively) supporting the notion that M and BS proteomes are deeply different. The functional categories of the differentially expressed proteins show that many metabolic processes are asymmetrically distributed between M and BS (Figure 2B), further supporting results obtained by previous maize and Setaria transcriptomic analyses (Chang *et al*., 2012, John *et al*., 2014). For example, protein homeostasis and modification, redox homeostasis, the oxidative pentose phosphate pathway, and lipid metabolism are enhanced in M, while more proteins related to Calvin cycle, photorespiration, Pyr oxidation, oxidative phosphorylation, and nucleotide metabolism are present in BS. In addition, functional categories such as light reactions, protein biosynthesis, amino acid metabolism, and solute transport show a similar number of proteins differentially accumulated in M and BS, emphasizing that those processes rely on the cooperation between both compartments (Figure 2B). The comparison of BS/M ratios from our data shows a Person’s correlation coefficient of 0.66 (*p*=4.884 E-68) with BS/M obtained by transcriptome analysis (John *et al*., 2014), which indicates a good agreement of protein and transcript levels (Supplementary Figure 2).

**Figure 2:**
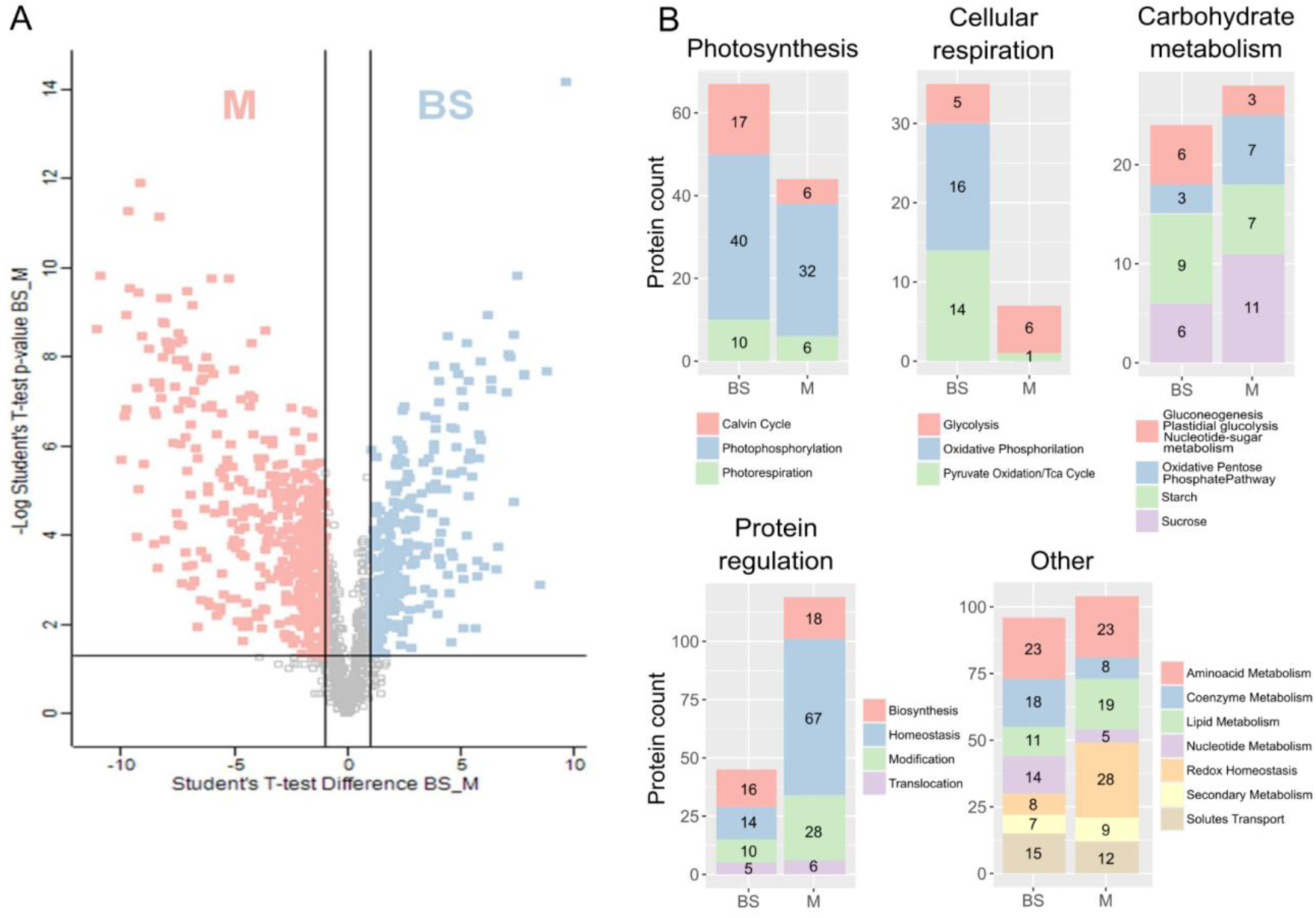
Protein distribution in M and BS from Setaria leaves. Volcano plot showing the proteins that were differentially enriched in M or in BS (light red). The horizontal line marks the limit of p-value<0.05, while the vertical lines mark the limit at which the difference in the proteins is more than doubled in one condition over the other **(A)**. Proteins identified according to their role, grouped by metabolism. Proteins significantly enriched in M or BS show that these cells fulfil different functions in metabolic and regulatory processes **(B)**.

Among the decarboxylases that can participate in the C_4_ cycle, three ME were detected in M and BS proteomes, one chloroplastic NADP-ME (C_4_-NADP-ME) and two mitochondrial NAD-ME (NAD-ME1 and NAD-ME2). No PEPCK peptide was detected, which is consistent with the low level of the transcript encoding this enzyme previously reported (John *et al*., 2014; de Oliveira Dal’Molin *et al*., 2016). Table 1 shows a comparison of the levels of these transcripts and proteins in maize and Setaria. The rest of the C_4_ cycle enzymes such as CA, PEPC and PPDK were more abundant in M samples in agreement with their photosynthetic roles (Supplementary Table 1).

**Table 1:** Convergence of NADP-ME and NAD-ME transcripts and proteins in Setaria viridis (Sv) and maize (Zm). Log_2_FC (BS/M) of maize transcripts and proteins were obtained from Chang *et al*., 2012 and Friso *et al*, 2010, respectively. Log_2_FC (BS/M) of Setaria transcripts and proteins were obtained from John *et al*., 2014 and this work, respectively. NA: not available.

|  | ID | Transcripts<br>(<sup>a</sup>John *et al.*, 2014; <sup>b</sup>Chang *et al.*, 2012) |  | Proteins<br>(<sup>c</sup>this work; <sup>d</sup>Friso *et al.*, 2010) |  |
| --- | --- | --- | --- | --- | --- |
|  |  | Log<sub>2</sub>FC(BS/M) | *p* adjusted | Log<sub>2</sub>FC(BS/M) | *p* adjusted |
| SvNAD-ME1 | Sevir.2G333400 | 0.7434<sup>a</sup> | 2.74E-03<sup>a</sup> | 0.8267<sup>c</sup> | 0.0284<sup>c</sup> |
| SvNAD-ME2 | Sevir.9G199800 | 0.8385<sup>a</sup> | 6.34E-04<sup>a</sup> | 0.8735<sup>c</sup> | 0.0593<sup>c</sup> |
| SvNADP-ME4 (C4) | Sevir.5G132500 | 7.2049<sup>a</sup> | 7.82E-84<sup>a</sup> | 5.4296<sup>c</sup> | 1.39E-06<sup>c</sup> |
| ZmNAD-ME1 | GRMZM2G085747 | 1.5706<sup>b</sup> | 5.46E-13<sup>b</sup> | NA | NA |
| ZmNAD-ME2 | GRMZM2G406672 | 3.4952<sup>b</sup> | 4.93E-38<sup>b</sup> | NA | NA |
| ZmNADP-ME (C4) | GRMZM2G085019 | 6.6616<sup>b</sup> | 1.20E-149<sup>b</sup> | 1.48<sup>d*</sup> | 9.13E-03<sup>d*</sup> |
| ZmNADP-ME (nonC4) | GRMZM2G122479 | NA | NA | 1.48<sup>d*</sup> | 9.13E-03<sup>d*</sup> |

\*were reported as a family group

### Chloroplastic NADP-ME is the main decarboxylase in C4 cycle and probably fulfils also non-photosynthetic roles in Setaria

The chloroplastic NADP-ME was highly represented in BS proteome, as expected, with an enrichment of 42-fold respect to M (Table 1). In the genomes of *S. viridis* and *S. italica* there is only one gene encoding a plastidic NADP-ME, which is identical in both species, therefore we will refer to them as Setaria C_4_-NADP-ME. The kinetic analysis of the purified recombinant enzyme shows that C_4_-NADP-ME of Setaria shares many characteristics with the maize and sorghum photosynthetic NADP-ME (Detarsio *et al*., 2003; Saigo *et al*., 2013a), such as high catalytic efficiency and Mal inhibition at pH 7.0 (Figure 3, Table 2). This can be attributed to the conserved amino acid residues important for the C_4_ role previously identified by crystallographic and site-directed mutagenesis analyses (Supplementary Figure 3, Alvarez *et al*., 2019). Similarly to maize C_4_-NADP-ME (Detarsio *et al*., 2007), the metabolites tested did not show significant effects on Setaria C_4_-NADP-ME activity and Mal inhibition was the main modulation. This response is important to prevent the overconsumption of Mal under dark conditions, when stromal pH decreases and is close to 7.0 (Saigo *et al*., 2013b). Considering that this is the only plastidic isoform encoded in the genome, it probably performs other roles in organs like roots and seeds and leaf M cells. In maize, the transcript encoding a second plastidic NADP-ME (nonC_4_-NADP-ME) has been detected in roots, grains, stems, and leaves where it provides Pyr and NADPH to non-photosynthetic pathways, like plastidic NADP-ME isoforms present in C_3_ plants (Gerrard Wheeler *et al*, 2005; Alvarez *et al*, 2013,). Consistent with this, Setaria C_4_-NADP-ME was detected in M protein samples (this study) and non-photosynthetic organs of Setaria (publicly available data, Supplementary Table 2). The protein sequence of Setaria C_4_-NADP-ME also shows residues conserved in nonC_4_-NADP-ME from maize, sorghum and rice (Supplementary Figure 3) and conserved regulatory elements were detected in these genes (Alvarez *et al*., 2013), which could be essential to fulfil non-photosynthetic roles.

**Figure 3:**
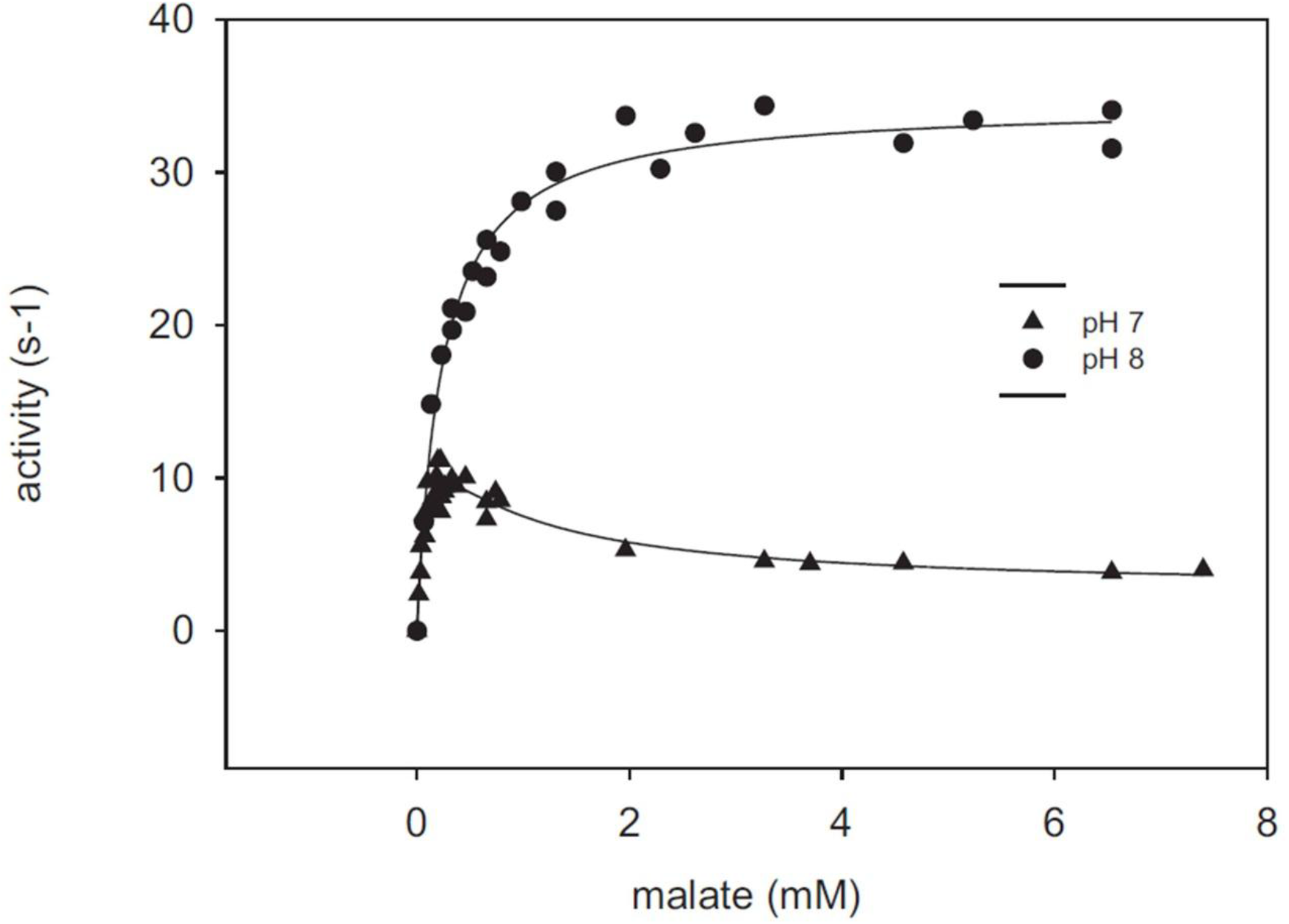
Kinetic and regulatory properties of Setaria C_4_-NADP-ME. NADP-ME activity was determined at different malate concentrations, at pH 8.0 and 7.0.

**Table 2:** Kinetic parameters of Setaria chloroplastic NADP-ME. Maize (Zm) and sorghum (Sb) C_4_ and nonC_4_ enzyme parameters were previously determined and included for comparison (Detarsio *et al*., 2003; Saigo *et al*., 2004; Detarsio *et al*., 2007; Saigo *et al*., 2013a). *k*cat corresponds to the μmol of substrate converted into product by s under optimal conditions per μmol of active site and S_0.5_ is the substrate concentration for which half of maximum reaction rate is obtained. %CV were below 5%.

|  | ZmnonC<sub>4</sub> | SbnonC<sub>4</sub> | SetariaC<sub>4</sub> | ZmC<sub>4</sub> | SbC<sub>4</sub> |
| --- | --- | --- | --- | --- | --- |
| Optimal pH | 8.0 | 8.0 | 8.0 | 8.0 | 8.0 |
| *k*<sub>cat</sub> (s<sup>-1</sup>) | 26.3 | 26.4 | 35.0 | 28.1 | 36.8 |
| S<sub>0.5</sub> NADP<sup>+</sup> (μM) | 70.2 | 16.8 | 9.3 | 8.0 | 11.0 |
| S<sub>0.5</sub> Mal (mM) | 0.8 | 0.6 | 0.18 | 0.2 | 0.4 |
| Inhibition by Mal at pH 7.0 | low | low | yes | yes | yes |

### Mitochondrial NAD-ME can collaborate in malate decarboxylation during photosynthesis

In plants, NAD-ME isoforms are exclusively mitochondrial and NADP-ME isoforms are present in cytosol and plastids. According to a 40% identity (on average), NAD- and NADP dependent enzymes share structural motifs, but they have arisen by different evolutionary events and evolved independently (Tronconi *et al*., 2018). All plant species conserve at least two NAD-dependent isoforms, one α-NAD-ME and one β-NAD-ME, which arose by a gene duplication that occurred late in the evolution of vascular plants (Tronconi *et al*., 2020). The α-NAD-ME and the β-NAD-ME encoded in the genome of *S. viridis*, also called NAD-ME1 and NAD-ME2, respectively, were detected in M and BS protein samples, but they are slightly more abundant in BS (FC(BS/M): 1.8 in both cases, Table 1). NAD-ME1 from *S. italica* and *S. viridis* are identical and the same occurs with NAD-ME2 proteins; then, they will be mentioned as Setaria NAD-ME1 and NAD-ME2. The kinetic parameters of the purified recombinant enzymes were similar to those reported for Arabidopsis isoforms (Table 3). In this C_3_ plant, NAD-ME1 and NAD-ME2 are hypothesized to participate in the nocturnal respiration, which is supported by a coordinated modulation of the activity by glycolytic and tricarboxylic acid (TCA) cycle intermediates (Tronconi *et al*., 2008 and 2010b). The percentage of identity of NAD-ME1 and NAD-ME2 of Setaria is high (65.8 %) and they are both strongly activated by citrate, α-ketoglutarate (α-KG), succinate, fumarate, fructose 1,6-bisphosphate (FBP) and PEP (Figure 4A and B). In addition, OAA, CoA and acetyl-CoA are positive modulators of NAD-ME2 (Figure 4B). These modulations indicate that NAD-ME1 and NAD-ME2 of Setaria could also be implicated in respiration. Nevertheless, unlike NADP-ME, the studies of photosynthetic NAD-ME are scarce and biochemical characteristics related to the photosynthetic role have not been described (Saigo *et al*., 2013b). Since Asp and alanine participate in the C_4_ carbon shuttle in NAD-ME subtype species and Glu mediates many transamination reactions, we evaluated if these metabolites modulate NAD-ME1 and NAD-ME2 activities. We could not detect any modulation by Asp and alanine, but Glu (2 mM) inhibited NAD-ME2 (Figure 4B). α-KG and Glu are intermediates in the nitrogen balancing reactions catalyzed by aminotransferases. In the reversible reaction catalyzed by aspartate aminotransferase (AAT), Asp is converted to OAA while α-KG is converted to Glu. Then, a high ratio of α-KG/Glu could enhance the production of OAA derived from Asp. After that, malate dehydrogenase (MDH) could reduce OAA to malate in a reversible reaction that is modulated by the availability of substrates and products. The reductive carboxylation of Pyr (reverse reaction) was not detected in our assay conditions with NAD-ME1 nor NAD-ME2, in agreement with Arabidopsis enzymes (Tronconi *et al*., 2010a; Badia *et al*., 2017). Therefore, unlike AAT and MDH, which catalyze reversible reactions, NAD-ME1 and NAD-ME2 represent regulatory spots that can have a great influence on organic acid metabolism. By this regulatory mechanism, NAD-ME2 activity and Mal availability can be readily coordinated by the ratio of α-KG/Glu (Figure 4C). Glu and α-KG are also connected by the Glu dehydrogenase (GDH) reaction. In our dataset we could detect that mitochondrial GDH1 and GDH2 were enriched in BS (FC(BS/M): 6.5 and 39.4, respectively; Table 4), which indicates that they could be influencing the ratio of α-KG/Glu in BS mitochondria. Although α-KG/Glu ratio could be a fine tuning of NAD-ME activity in the context of C_4_ cycle, more evidence is needed to prove this hypothesis. Together, the partial enrichment of NAD-ME1 and NAD-ME2 in BS, and their regulatory properties suggest that these proteins could participate in the decarboxylation of Mal in the mitochondria in cooperation with chloroplastic NADP-ME. Direct evidence of a species using both NADP-ME and NAD-ME for photosynthesis is limited, but recent works have noted elevated transcript of both decarboxylases within the same species (Rao *et al*., 2016; Washburn *et al*. 2017). In addition, a multi-omics analysis of *S. italica* young and mature leaves, provided more evidence suggesting that this species could use a mixed C_4_ decarboxylation mode that involves NADP-ME and NAD-ME (de Oliveira Dal’Molin *et al*., 2016).

**Table 3:** Comparative summary of the most important biochemical characteristics of Setaria NAD-MEs. Arabidopsis (At) NAD-ME parameters (Tronconi *et al*., 2008) were included for comparison. *k*cat corresponds to the μmol of substrate converted into product by s under optimal conditions per μmol of active site and S0.5 is the substrate concentration for which half of maximum reaction rate is obtained. %CV were below 5%.

|  | SetariaNAD-ME1 | SetariaNAD-ME2 | AtNAD-ME1 | AtNAD-ME2 |
| --- | --- | --- | --- | --- |
| Optimal pH | 6.5 | 6.5 | 6.4 | 6.6 |
| $k_{cat}$ ($s^{-1}$) | 12.7 | 15.5 | 31.1 | 44.1 |
| $S_{0.5}$ NAD<sup>+</sup> (mM) | 1.8 | 0.5 | 0.5 | 0.5 |
| $S_{0.5}$ Mal (mM) | 3.9 | 3.4 | 3.0 | 3.0 |

**Figure 4:**
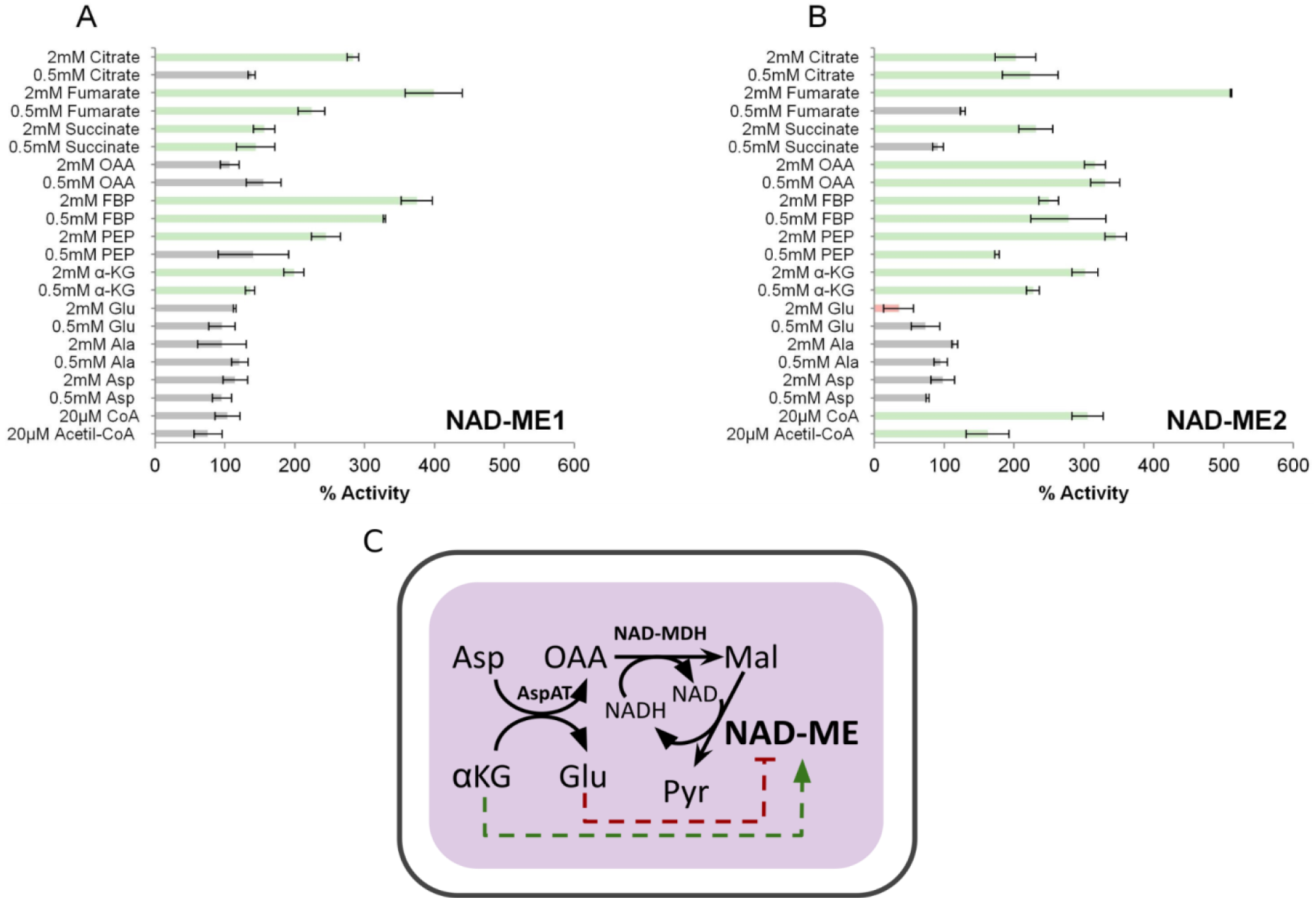
Regulatory properties of recombinant NAD-ME1 (A) and NAD-ME2 (B), The results represent the % of activity in the presence of each effector in relation to the activity measured in the absence of the metabolites (100%). Assays were performed by triplicate, and error bars indicate S.D. Bars in red, inhibition (less than 70% residual activity). Bars in green, activation (more than 140%).Schematic representation showing that NAD-ME activity and Mal availability can be readily coordinated by the ratio of α-KG/Glu **(C)**.Green dashed line indicates activation and red dashed line indicates inhibition.

**Table 4:**
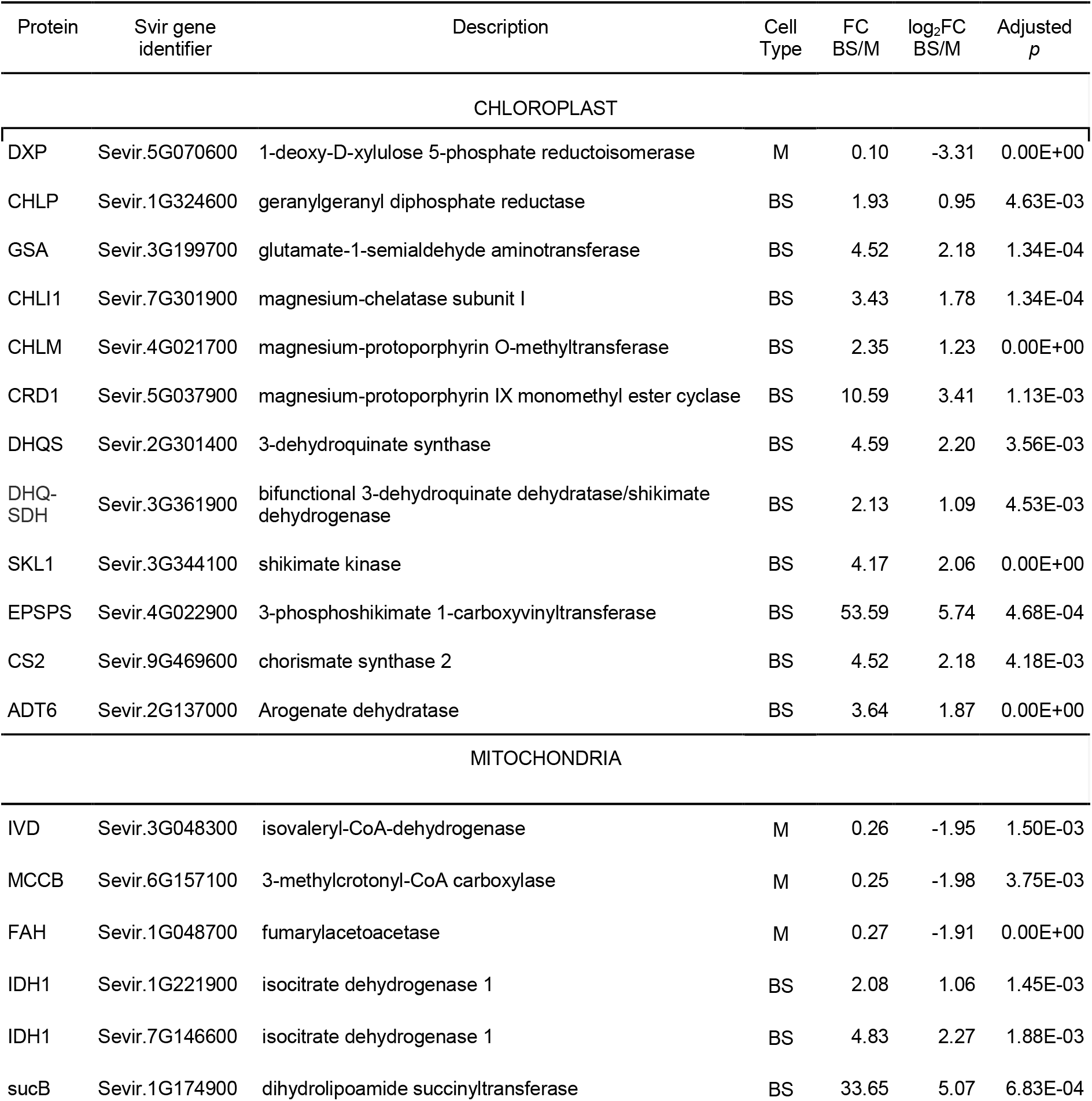
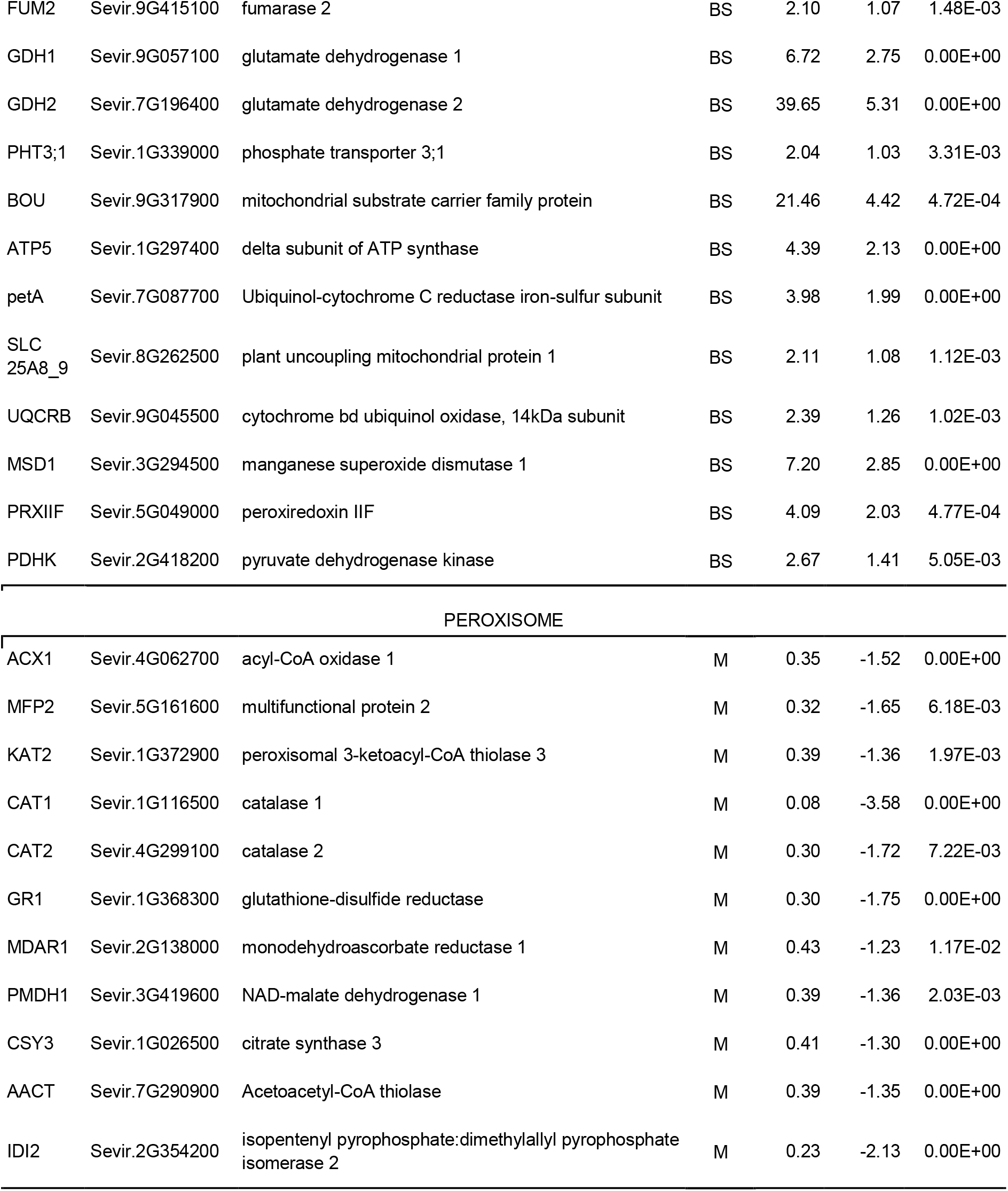
Experimental data of proteins enriched in M or BS. The putative subcellular location of each protein was inferred by the identification of its ortholog in proteomic databases of maize chloroplasts (Friso et al., 2010), Arabidopsis mitochondria (Fuchs et al., 2020) and Arabidopsis peroxisomes (Pan and Hu, 2018). The chloroplastic or mitochondrial location was also predicted by TargetP.

| Protein | Svir gene identifier | Description | Cell Type | FC BS/M | log<sub>2</sub>FC BS/M | Adjusted $p$ |
| --- | --- | --- | --- | --- | --- | --- |
| CHLOROPLAST |  |  |  |  |  |  |
| DXP | Sevir.5G070600 | 1-deoxy-D-xylulose 5-phosphate reductoisomerase | M | 0.10 | -3.31 | 0.00E+00 |
| CHLP | Sevir.1G324600 | geranylgeranyl diphosphate reductase | BS | 1.93 | 0.95 | 4.63E-03 |
| GSA | Sevir.3G199700 | glutamate-1-semialdehyde aminotransferase | BS | 4.52 | 2.18 | 1.34E-04 |
| CHLI1 | Sevir.7G301900 | magnesium-chelatase subunit I | BS | 3.43 | 1.78 | 1.34E-04 |
| CHLM | Sevir.4G021700 | magnesium-protoporphyrin O-methyltransferase | BS | 2.35 | 1.23 | 0.00E+00 |
| CRD1 | Sevir.5G037900 | magnesium-protoporphyrin IX monomethyl ester cyclase | BS | 10.59 | 3.41 | 1.13E-03 |
| DHQS | Sevir.2G301400 | 3-dehydroquinate synthase | BS | 4.59 | 2.20 | 3.56E-03 |
| DHQ-SDH | Sevir.3G361900 | bifunctional 3-dehydroquinate dehydratase/shikimate dehydrogenase | BS | 2.13 | 1.09 | 4.53E-03 |
| SKL1 | Sevir.3G344100 | shikimate kinase | BS | 4.17 | 2.06 | 0.00E+00 |
| EPSPS | Sevir.4G022900 | 3-phosphoshikimate 1-carboxyvinyltransferase | BS | 53.59 | 5.74 | 4.68E-04 |
| CS2 | Sevir.9G469600 | chorismate synthase 2 | BS | 4.52 | 2.18 | 4.18E-03 |
| ADT6 | Sevir.2G137000 | Arogenate dehydratase | BS | 3.64 | 1.87 | 0.00E+00 |
| MITOCHONDRIA |  |  |  |  |  |  |
| IVD | Sevir.3G048300 | isovaleryl-CoA-dehydrogenase | M | 0.26 | -1.95 | 1.50E-03 |
| MCCB | Sevir.6G157100 | 3-methylcrotonyl-CoA carboxylase | M | 0.25 | -1.98 | 3.75E-03 |
| FAH | Sevir.1G048700 | fumarylacetoacetase | M | 0.27 | -1.91 | 0.00E+00 |
| IDH1 | Sevir.1G221900 | isocitrate dehydrogenase 1 | BS | 2.08 | 1.06 | 1.45E-03 |
| IDH1 | Sevir.7G146600 | isocitrate dehydrogenase 1 | BS | 4.83 | 2.27 | 1.88E-03 |
| sucB | Sevir.1G174900 | dihydrolipoamide succinyltransferase | BS | 33.65 | 5.07 | 6.83E-04 |
| FUM2 | Sevir.9G415100 | fumarase 2 | BS | 2.10 | 1.07 | 1.48E-03 |
| GDH1 | Sevir.9G057100 | glutamate dehydrogenase 1 | BS | 6.72 | 2.75 | 0.00E+00 |
| GDH2 | Sevir.7G196400 | glutamate dehydrogenase 2 | BS | 39.65 | 5.31 | 0.00E+00 |
| PHT3;1 | Sevir.1G339000 | phosphate transporter 3;1 | BS | 2.04 | 1.03 | 3.31E-03 |
| BOU | Sevir.9G317900 | mitochondrial substrate carrier family protein | BS | 21.46 | 4.42 | 4.72E-04 |
| ATP5 | Sevir.1G297400 | delta subunit of ATP synthase | BS | 4.39 | 2.13 | 0.00E+00 |
| petA | Sevir.7G087700 | Ubiquinol-cytochrome C reductase iron-sulfur subunit | BS | 3.98 | 1.99 | 0.00E+00 |
| SLC<br>25A8_9 | Sevir.8G262500 | plant uncoupling mitochondrial protein 1 | BS | 2.11 | 1.08 | 1.12E-03 |
| UQCRB | Sevir.9G045500 | cytochrome bd ubiquinol oxidase, 14kDa subunit | BS | 2.39 | 1.26 | 1.02E-03 |
| MSD1 | Sevir.3G294500 | manganese superoxide dismutase 1 | BS | 7.20 | 2.85 | 0.00E+00 |
| PRXIIF | Sevir.5G049000 | peroxiredoxin IIF | BS | 4.09 | 2.03 | 4.77E-04 |
| PDHK | Sevir.2G418200 | pyruvate dehydrogenase kinase | BS | 2.67 | 1.41 | 5.05E-03 |
| PEROXISOME |  |  |  |  |  |  |
| ACX1 | Sevir.4G062700 | acyl-CoA oxidase 1 | M | 0.35 | -1.52 | 0.00E+00 |
| MFP2 | Sevir.5G161600 | multifunctional protein 2 | M | 0.32 | -1.65 | 6.18E-03 |
| KAT2 | Sevir.1G372900 | peroxisomal 3-ketoacyl-CoA thiolase 3 | M | 0.39 | -1.36 | 1.97E-03 |
| CAT1 | Sevir.1G116500 | catalase 1 | M | 0.08 | -3.58 | 0.00E+00 |
| CAT2 | Sevir.4G299100 | catalase 2 | M | 0.30 | -1.72 | 7.22E-03 |
| GR1 | Sevir.1G368300 | glutathione-disulfide reductase | M | 0.30 | -1.75 | 0.00E+00 |
| MDAR1 | Sevir.2G138000 | monodehydroascorbate reductase 1 | M | 0.43 | -1.23 | 1.17E-02 |
| PMDH1 | Sevir.3G419600 | NAD-malate dehydrogenase 1 | M | 0.39 | -1.36 | 2.03E-03 |
| CSY3 | Sevir.1G026500 | citrate synthase 3 | M | 0.41 | -1.30 | 0.00E+00 |
| AACT | Sevir.7G290900 | Acetoacetyl-CoA thiolase | M | 0.39 | -1.35 | 0.00E+00 |
| IDI2 | Sevir.2G354200 | isopentenyl pyrophosphate:dimethylallyl pyrophosphate isomerase 2 | M | 0.23 | -2.13 | 0.00E+00 |

### Chloroplastic and mitochondrial proteins sustain different metabolisms in M and BS

In order to organize the data and to obtain a clear view, we sub-classified the set of proteins enriched in M or BS (log_2_FC(BS/M)<−1 or >1, *p*-value<0.05) according to their putative location in the cell. We utilized a combination of targeting signals detection by bioinformatic tools and the identification of Setaria orthologues of organellar proteins previously detected by proteomic approaches (Friso *et al*., 2010, Fuchs *et al*., 2020). This strategy helped us to visualize the main differences in the metabolic demands of M and BS in Setaria. We assigned 142 chloroplastic proteins in M and 177 in BS, and 25 mitochondrial proteins in M and 30 in BS (Supplementary Table 1).

In C_4_ species, BS chloroplasts hold much of the photosynthetic machinery since most carbon assimilation steps operate almost exclusively in these organelles. In the NADP-ME subtype, the high amount and catalytic proficiency of chloroplastic NADP-ME provides CO_2_ and NADPH to the Calvin cycle. The ATP is generated by the cyclic electron transport around PSI and the trioseP/PGA shuttle contributes to the NADPH needed for carbon assimilation. The chloroplastic proteomes of M and BS in maize provided strong experimental support for a number of specialized BS and M metabolic functions, including starch biosynthesis (BS), methylerythritol phosphate (MEP) pathway for isoprenoid biosynthesis (M), nitrogen assimilation (M), the initial steps in sulfur assimilation (BS), among others (Friso *et al*., 2010). Here, we obtained evidence that shows that many of the M and BS specific metabolic functions are conserved in Setaria chloroplasts, but some others are different from maize (Supplementary Table 1). The comparison of the distribution and level of chloroplastic proteins of maize and *S. viridis* shows a Pearson correlation coefficient of 0.28 (*p*=6.22 E-10, Supplementary Figure 4). Among the 452 proteins included in the analysis (chloroplastic, *p*<0.05), 254 were detected in the same sample compartment (M or BS) in both species. For example, 1-deoxy-D-xylulose 5-phosphate reductase (DXR), the second enzyme of the MEP pathway, accumulated at higher levels in the M than in BS chloroplasts (Table 4), which can be attributed to a high demand of tocopherol to protect membranes from oxidative damage derived from the linear electron transport in M chloroplasts (Soll *et al*., 1983; Lichtenthaler *et al*., 1997). On the other hand, 165 maize M-enriched proteins were identified as BS-enriched in Setaria. For example, many enzymes involved in chlorophyll biosynthesis (glutamate-1-semialdehyde-2,1-aminomutase, Mg-protoporphyrin IX chelatase, Mg-protoporphyrin O-methyltransferase and Mg-protoporphyrin IX monomethyl ester cyclase) and the shikimate pathway for aromatic amino acids biosynthesis (3-dehydroquinate synthase, 3-dehydroquinate dehydratase, shikimate kinase and 5-enolpyruvyl-shikimate 3-phosphate synthase) are enhanced in Setaria BS chloroplasts (Table 4), but these pathways are enriched in maize M chloroplasts (Friso *et al*., 2010). Chorismate synthase and arogenate dehydratase, the enzymes which catalyze the last steps in chorismate and phenylalanine biosynthesis respectively, are also upregulated in BS (Table 4), but their presumably chloroplastic location could not be verified in this work. The weak correlation of M and BS chloroplast proteins of maize and *S. viridis* was already noted when comparing Setaria RNA-Seq and maize proteomics (John *et al*., 2014). These observations could be attributed to species-related divergences and could also point to differences between leaf tip (Friso *et al*., 2010) and middle blade (this study). Detailed studies of the proteomic, transcriptomic and metabolomic patterns along the maize leaf have shown that the metabolism largely varies from the heterotrophic (sink) base to the photosynthetic (source) tip (Majeran *et al*., 2010; Pick *et al*., 2011). While the region nearer to the base is characterized by the biosynthesis of lipids, cell wall, lignins and isoprenoids (MVA pathway) to support the expansion of the cells, the tip is engaged in the highly efficient photosynthetic C_4_ cycle to provide carbon compounds to other tissues. This distribution of roles helps to avoid the interference between photosynthetic and other metabolisms. In this work, we analyzed the central portion of *S. viridis* leaf where other strategies should be used to maintain the metabolic homeostasis of the cells. In this sense, a lower activity of the shikimate pathway in M would prevent the consumption of PEP, thus avoiding an interference with the C_4_ cycle.

A small group of mitochondrial proteins were significantly enriched in M or BS. In M, enzymes involved in the degradation of leucine (isovaleryl-CoA-dehydrogenase, 3-methylcrotonyl-CoA carboxylase) and phenylalanine (fumarylacetoacetate hydrolase) seems to be enhanced (Table 4). Despite the catabolism of both amino acids are completely different, both contribute to the pool of Acetyl-CoA. In BS mitochondria, there are higher levels of TCA cycle and TCA cycle-related enzymes (isocitrate dehydrogenase 1, dihydrolipoamide succinyltransferase, fumarase 2, Glu dehydrogenase 1 and 2), phosphate and carbon compound transporters (phosphate transporter 3;1, mitochondrial substrate carrier family protein), electron transport chain and oxidative phosphorylation components (delta subunit of ATP synthase, ubiquinol-cytochrome C reductase iron-sulfur subunit, plant uncoupling mitochondrial protein 1, cytochrome bd ubiquinol oxidase 14 kDa subunit) and redox response proteins (manganese superoxide dismutase 1, peroxiredoxin IIF), which indicates an active oxidative metabolism (Table 4). However, the accumulation of pyruvate dehydrogenase kinase (PDHK, Table 4) in this organelle entails that pyruvate dehydrogenase complex could be inhibited, opening the question of which is the substrate to be oxidized by TCA cycle. In this regard, it is important to recall that in plants the enzymatic activities of the TCA cycle can operate in diverse metabolic contexts in non-cyclic fashions, resembling more to an organic acids network deeply connected with the cytosol than to a close mitochondrial cycle (Sweetlove *et al*., 2010). The enrichment of PDHK in mitochondria of BS has been pointed out as a NAD-ME C_4_ pathway marker, since pyruvate generated by NAD-ME needs to be preserved to replenish PEP in M chloroplasts (Bräutigam *et al*, 2014). This observation also reinforces the idea that in Setaria NAD-ME activity in BS mitochondria could be actively collaborating in the C_4_ metabolism. Considering that in BS chloroplasts pyruvate consuming pathways could be active, such as beta-reduction for fatty acids and isoprenoids biosynthesis, at least at a low rate, the decarboxylation of malate in the mitochondria would provide an alternative route to preserve pyruvate for photosynthetic PEP recycling. Avoiding the leakage of intermediates from the C_4_ core to other metabolic routes is critical for the robustness of the C_4_ cycle functioning (Bräutigam *et al*, 2014). In this work we present evidence of several strategies to uncouple the C_4_ cycle from other pathways.

### Peroxisomal production of Acetyl-CoA is enhanced in M

In C_3_ plants,peroxisomes collaborates with chloroplasts and mitochondria in the recycling of PG carbon by photorespiration among many other metabolic functions such as fatty acid β-oxidation and phytohormone biosynthesis (see Kao *et al*., 2018 for a review). In C_4_ plants photorespiration has been minimized and restricted to BS, the site where PG is produced. To further investigate the metabolic routes that operate in peroxisomes of M and BS, we identified a group of putative peroxisomal proteins by retrieving the orthologues of Arabidopsis peroxisomal proteins from the set of proteins enriched in Setaria M or BS (Pan and Hu, 2018). In the M-enriched set, 29 peroxisomal proteins were identified and 2 proteins were identified in the BS-enriched group (Supplementary Table 1). The functional categories of M proteins show an active Acetyl-CoA production by β-oxidation of fatty acids (acyl-CoA oxidase 1, 3-hydroxyacyl-CoA dehydrogenase, 3-ketoacyl-CoA thiolase 3), which would be the main source of H_2_O_2_ instead of photorespiration (Table 4). This elevated production of H_2_O_2_ can be compensated by the upregulation of catalases 1 and 2 and ascorbate-glutathione redox system (glutathione reductase, monodehydroascorbate reductase 1) to protect the cell from oxidative damage (Table 4). Acetyl-CoA thiolase and isopentenyl diphosphate (IPP) isomerase accumulated at higher levels in M peroxisomes than in BS (Table 4) thus potentially providing a higher level of IPP and dimethylallyl pyrophosphate (DMAPP), to feed biosynthesis of terpenoids and phytosterols through the mevalonate (MVA) pathway (Vranová *et al*., 2013). The produced Acetyl-CoA probably does not fuel gluconeogenesis, since the glyoxylate pathway diminishes as seedlings mature (Titus and Becker, 1985) and Mal synthase was not detected in our data. The accumulation of malate dehydrogenase (MDH) and citrate synthase in peroxisomes (Table 4) and ATP-citrate lyase in the cytosol of M (log_2_FC(BS/M): −1.54; Supplementary Table 1) could be supporting the active export of Acetyl-CoA by the citrate shuttle (Table 4). In the cytosol, Acetyl-CoA is able to feed other biosynthetic routes. For example, the acetylation of serine using Acetyl-CoA is the first step to cysteine biosynthesis, which could be enhanced in M since the second enzyme is enriched in this compartment (O-acetylserine (thiol) lyase, Sevir.3G415300, log_2_FC(BS/M): −2.5; Supplementary Table 1). The Acetyl-CoA could be further incorporated in the nucleus to support the acetylation of histones. In this sense, it was recently found that peroxisomal β-oxidation regulates histone acetylation and DNA methylation in Arabidopsis (Wang *et al*., 2019) and the acetylation of histones in PEPC and ME promoters modulates the expression of these genes (Heinmann *et al*., 2013), which opens the question of the relation of peroxisomal metabolism and epigenetics in defining the expression patterns in M and BS.

### Concluding remarks

In this work we present evidence of a primary and a secondary route of Mal decarboxylation in BS cells (Figure 5). This would allow the plant to maintain the photosynthetic performance even under fluctuating environmental conditions. Future work is needed to better describe this auxiliary response and its contribution under stress conditions.

**Figure 5:**
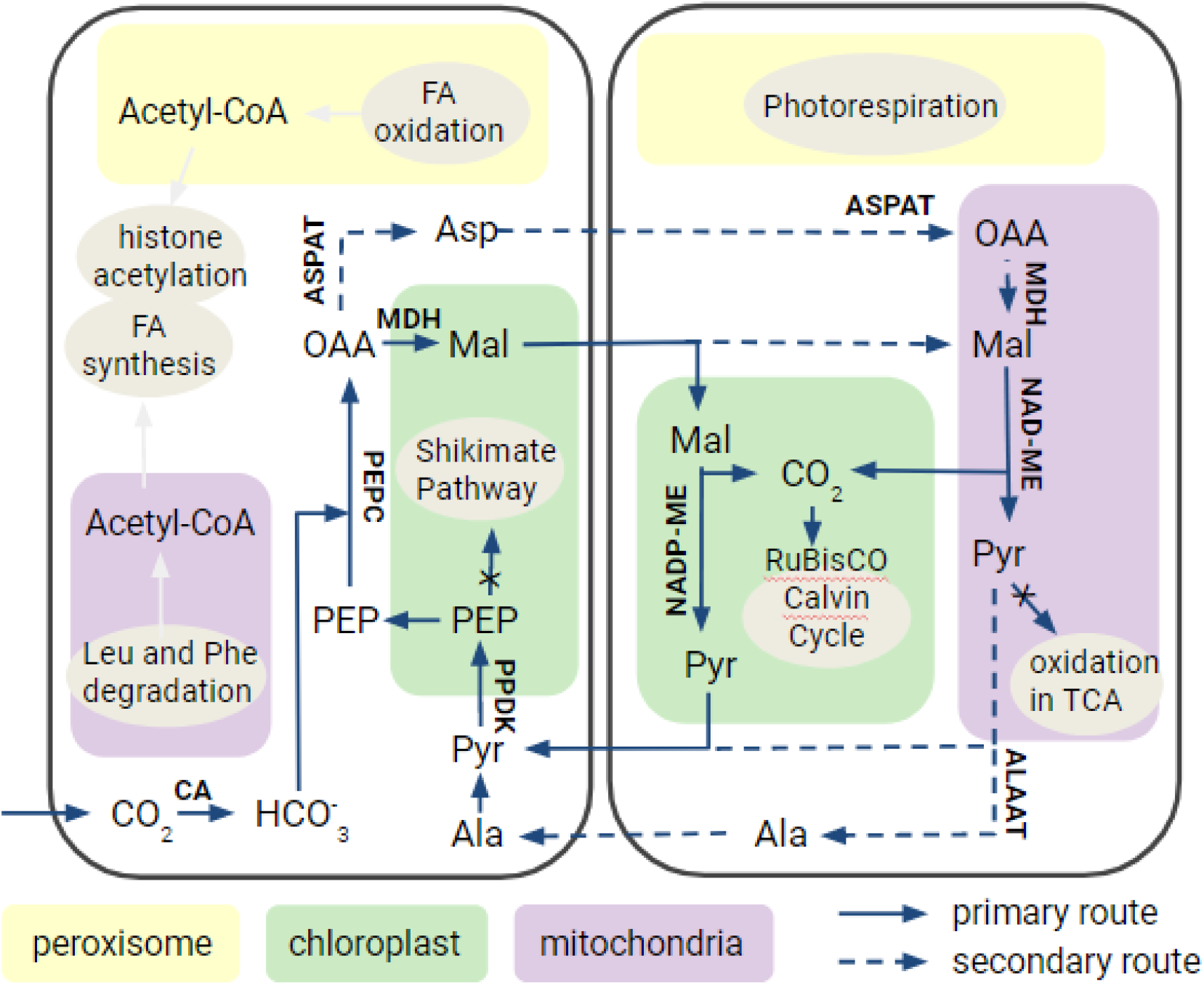
Schematic model of the C_4_ cycle in *S. viridis*. Besides the primary route of Mal decarboxylation in chloroplasts of BS cells, a secondary route in the mitochondria could also contribute to the C_4_ carbon shuttle. The pyruvate generated in the mitochondria of BS would not be oxidized to form Acetyl-CoA, while in M several Pyr-independent pathways of Acetyl-CoA generation are enhanced. In both cases, the Pyr pool would be preserved to maintain the C_4_ shuttle. Additionally, in M chloroplasts the down-regulation of the Shikimate pathway would reserve the PEP for the primary carboxylation by PEPC.

Based on comparative proteomics of M and BS, we found evidence of strategies to avoid the leakage of C_4_ intermediates. The pyruvate generated in the mitochondria of BS would not be oxidized to form acetyl-CoA, while in M several Pyr-independent pathways of acetyl-CoA generation are enhanced. In both cases, the Pyr pool would be preserved to maintain the C_4_ shuttle. Additionally, in M chloroplasts the down-regulation of the Shikimate pathway would reserve the PEP for the primary carboxylation by PEPC (Figure 5).

The C_4_ leaf presents transversal (M-BS) and longitudinal (base-tip) metabolic gradients that should be finely coordinated to achieve high rates of carbon assimilation. In those segments where there is an overlapping of several pathways, a fine regulation is needed to maintain the metabolic homeostasis. Although a great amount of multi-omics studies are facing this subject, more research on post-translational modification and enzymatic regulation are needed.

## Abbreviations

3PG 3: phosphoglycerate
AAT: aspartate aminotransferase BS bundle sheath
CA: carbonic anhydrase
CCM: carbon concentration mechanism FBP fructose 1,6-bisphosphate
GDH: glutamate dehydrogenase M mesophyll
Mal: malate
MDH: malate dehydrogenase MEP Methylerythritol Phosphate MVA mevalonic acid
NAD-ME: NAD-malic enzyme NADP-ME NADP-malic enzyme OAA oxaloacetate
Os: *Oryza sativa*
PDHK: pyruvate dehydrogenase kinase PEP phosphoenolpyruvate
PEPC: PEP carboxylase
PEPCK: phosphoenolpyruvate carboxylase PG 2-phosphoglycolate
PPDK: pyruvate phosphate dikinase Pyr pyruvate
RuBP: ribulose-1,5-bisphosphate Sb *Sorhum bicolor*
Si: *Setaria itálica*
Sv: *Setaria viridis* TCA tricarboxylic acid Zm *Zea mays*
α-KG: α-ketoglutarate

## Supplementary data

Supplementary Figure 1: SDS-PAGE of total protein extraction of M and BS.

Supplementary Figure 2: Correlation of transcripts and proteins in Setaria leaves.

Supplementary Figure 3: Amino acid residues conserved in C_4_ and nonC_4_ isoforms of NADP-ME from Setaria (Si), maize (Zm), sorghum (Sb) and rice (Os).

Supplementary Figure 4: Correlation of Setaria and maize orthologs.

Supplementary Table 1: Proteomic data and analysis.

Supplementary Table 2: Expression of Setaria C_4_-NADP-ME in various tissues and organs. Source: Thomas Brutnell, Danford Center (https://phytozome.jgi.doe.gov/phytomine/portal.do?externalid=PAC:32682171&class=gene)

## Acknowledgements

EM, CMF, CSA, MCGW, and MS belong to the Researcher Career of National Council of Scientific and Technical Research (CONICET). PC and TT are fellows of the same institution. This work was financially supported by the National Agency for Promotion of Science and Technology grants to CSA (PICT-2014-2556) and MS (PICT-2019-02890). CMF is funded by the National Agency for Promotion of Science and Technology (PICT-2018-00865) and the Max Planck Society (Partner Group for Plant Biochemistry).

## Author contributions

CMF, CSA, MCGW, and MS designed the concept and planned the experiments of this study. PC, TT, CMF, and MS performed the experiments regarding M and BS separation, protein extraction, and sample preparation for mass spectrometry. PC, EM, and MS were involved in the bioinformatics analysis. PC, CL, MCGW, and MS carried out the cloning, expression, and kinetic characterization of the recombinant enzymes. PC, EM, CMF, MCGW, and MS were involved in the analysis and interpretation of the results. MS and PC, in collaboration with all the other authors, wrote and edited the manuscript.

## Data Availability Statement

All data supporting the findings of this study are available within the paper and within its supplementary materials published online.

## Notes

### Competing Interest Statement

The authors have declared no competing interest.

## References

Acharya BR, Roy Choudhury S, Estelle AB, Vijayakumar A, Zhu C, Hovis L, Pandey S (2017) Optimization of phenotyping assays for the model monocot *Setaria viridis*. Front. Plant Sci. 8: 2172.

Almagro Armenteros JJ, Salvatore M, Emanuelsson O, Winther O, von Heijne G, Elofsson A, Nielsen H (2019) Detecting sequence signals in targeting peptides using deep learning. Life Sci. Alliance 2: e201900429.

Alvarez CE, Bovdilova A, Höppner A, Wolff CC, Saigo M, Trajtenberg F, Zhang T, Buschiazzo A, Nagel-Steger L, Drincovich MF, Lercher MJ, Maurino VG (2019) Molecular adaptations of NADP-malic enzyme for its function in C_4_ photosynthesis in grasses. Nat. Plants 5: 755–765.

Alvarez CE, Saigo M, Margarit E, Andreo CS, Drincovich MF (2013) Kinetics and functional diversity among the five members of the NADP-malic enzyme family from. Photosynth. Res. 115: 65–80.

Badia MB, Mans R, Lis AV, Tronconi MA, Arias CL, Maurino VG, Andreo CS, Drincovich MF, van Maris AJ, Gerrard Wheeler MC (2017) Specific *Arabidopsis thaliana* malic enzyme isoforms can provide anaplerotic pyruvate carboxylation function in *Saccharomyces cerevisiae*. FEBS J. 284: 654–665.

Bailey-Serres J, Parker JE, Ainsworth EA, Oldroyd GED, Schroeder JI (2019) Genetic strategies for improving crop yields. Nature 575: 109–118.

Bauwe H, Hagemann M, Fernie AR (2010) Photorespiration: players, partners and origin. Trends Plant Sci. 15: 330–336.

Bennetzen JL, Schmutz J, Wang H, Percifield R, Hawkins J, Pontaroli AC, Estep M, Feng L, Vaughn JN, Grimwood J, Jenkins J, Barry K, Lindquist E, Hellsten U, Deshpande S, Wang X, Wu X, Mitros T, Triplett J, Yang X, Ye CY, Mauro-Herrera M, Wang L, Li P, Sharma M, Sharma R, Ronald PC, Panaud O, Kellogg EA, Brutnell TP, Doust AN, Tuskan GA, Rokhsar D, Devos KM (2012) Reference genome sequence of the model plant Setaria. Nat. Biotechnol. 30: 555–561.

Bradford MM (1976) A rapid and sensitive method for the quantitation of microgram quantities of protein utilizing the principle of protein-dye binding.Anal Biochem. 72: 248–254.

Bräutigam A, Schliesky S, Külahoglu C, Osborne CP, Weber APM (2014) Towards an integrative model of C_4_ photosynthetic subtypes: insights from comparative transcriptome analysis of NAD-ME, NADP-ME, and PEP-CK C_4_ species. J. Exp. Bot. 65: 3579–3593.

Burnette WN (1981) “Western blotting”: electrophoretic transfer of proteins from sodium dodecyl sulfate-polyacrylamide gels to unmodified nitrocellulose and radiographic detection with antibody and radioiodinated protein A. Anal. Biochem. 112: 195–203.

Covshoff S, Furbank RT, Leegood RC, Hibberd JM (2013) Leaf rolling allows quantification of mRNA abundance in mesophyll cells of sorghum. J. Exp. Bot. 64: 807–813.

de Oliveira Dal’Molin CG, Orellana C, Gebbie L, Steen J, Hodson MP, Chrysanthopoulos P, Plan MR, McQualter R, Palfreyman RW, Nielsen LK (2016) Metabolic reconstruction of *Setaria italica*: A systems biology approach for integrating tissue-specific omics and pathway analysis of bioenergy grasses. Front. Plant Sci. 7: 1138.

Detarsio E, Alvarez CE, Saigo M, Andreo CS, Drincovich MF (2007) Identification of domains involved in tetramerization and malate inhibition of maize C_4_-NADP-malic enzyme. J. Biol. Chem. 282: 6053–6060.

Detarsio E, Wheeler MC, Campos Bermúdez VA, Andreo CS, Drincovich MF (2003) Maize C_4_ NADP-malic enzyme. Expression in *Escherichia coli* and characterization of site-directed mutants at the putative nucleoside-binding sites. J. Biol. Chem. 278: 13757–13764.

Drincovich MF, Lara MV, Maurino VG, Andreo CS (2011) C_4_ decarboxylases: different solutions for the same biochemical problem, the provision of CO_2_ to Rubisco in the bundle sheath cells. In: C_4_ photosynthesis and related CO_2_ concentrating mechanisms. Raghavendra AS, Sage RF (eds.), Springer Netherlands: 277–300.

Doust AN, Brutnell TP, Upadhyaya HD, Van Eck J (2019) Editorial: *Setaria* as a model genetic system to accelerate yield increases in cereals, forage crops, and bioenergy grasses. Front. Plant Sci. 10: 1211.

Edwards EJ, Smith SA (2010) Phylogenetic analyses reveal the shady history of C_4_ grasses. Proc. Natl. Acad. Sci. USA 107: 2532–2537.

Friso G, Majeran W, Huang M, Sun Q, van Wijk KJ (2010). Reconstruction of metabolic pathways, protein expression, and homeostasis machineries across maize bundle sheath and mesophyll chloroplasts: large-scale quantitative proteomics using the first maize genome assembly. Plant Physiol. 152: 1219–1250.

Fuchs P, Rugen N, Carrie C, Elsässer M, Finkemeier I, Giese J, Hildebrandt TM, Kühn K, Maurino VG, Ruberti C, Schallenberg-Rüdinger M, Steinbeck J, Braun HP, Eubel H, Meyer EH, Müller-Schüssele SJ, Schwarzländer M (2020) Single organelle function and organization as estimated from Arabidopsis mitochondrial proteomics. Plant J. 101: 420–441.

Furbank RT (2011) Evolution of the C_4_ photosynthetic mechanism: are there really three C_4_ acid decarboxylation types? J. Exp. Bot. 62: 3103–3108.

Ghannoum O, Evans JR, Caemmerer S (2011) Nitrogen and water use efficiency of C_4_ plants. In: C_4_ photosynthesis and related CO_2_ concentrating mechanisms. Raghavendra AS, Sage RF (eds.), Springer Netherlands: 129–146.

Gerrard Wheeler MC, Tronconi MA, Drincovich MF, Andreo CS, Flügge UI, Maurino VG (2005) A comprehensive analysis of the NADP-malic enzyme gene family of Arabidopsis. Plant Physiol. 139: 39–51.

Grover SD, Canellas PF, Wedding RT (1981) Purification of NAD malic enzyme from potato and investigation of some physical and kinetic properties. Arch. Biochem. Biophys. 209: 396–407.

Hatch MD (1987) C_4_ photosynthesis: a unique blend of modified biochemistry, anatomy and ultrastructure. Biochim.Biophys. Acta 895: 81–106.

Heimann L, Horst I, Perduns R, Dreesen B, Offermann S, Peterhänsel C (2013) A common histone modification code on C_4_ genes in maize and its conservation in *Sorghum bicolor* and *Setaria italica*. Plant Physiol. 162: 456–469.

Howe KL, Contreras-Moreira B, De Silva N, Maslen G, Akanni W, Allen J, Alvarez-Jarreta J, Barba M, Bolser DM, Cambell L, Carbajo M, Chakiachvili M, Christensen M, Cummins C, Cuzick A, Davis P, Fexova S, Gall A, George N, Gil L, Gupta P, Hammond-Kosack KE, Haskell E, Hunt SE, Jaiswal P, Janacek SH, Kersey PJ, Langridge N, Maheswari U, Maurel T, McDowall MD, Moore B, Muffato M, Naamati G, Naithani S, Olson A, Papatheodorou I, Patricio M, Paulini M, Pedro H, Perry E, Preece J, Rosello M, Russell M, Sitnik V, Staines DM, Stein J, Tello-Ruiz MK, Trevanion SJ, Urban M, Wei S, Ware D, Williams G, Yates AD, Flicek P (2020) Ensembl Genomes 2020-enabling non-vertebrate genomic research. Nucleic Acids Res. 48, D689–D695.

John CR, Smith-Unna RD, Woodfield H, Covshoff S, Hibberd JM (2014). Evolutionary convergence of cell-specific gene expression in independent lineages of C_4_ grasses. Plant Physiol. 165: 62–75.

Jordan DB, Ogren WL (1984) The CO_2_/O_2_ specificity of ribulose 1,5-bisphosphate carboxylase/oxygenase. Planta 161: 308–313.

Kao YT, Gonzalez KL, Bartel B (2018) Peroxisome function, biogenesis, and dynamics in plants. Plant Physiol. 176: 162–177.

Kinsella RJ, Kähäri A, Haider S, Zamora J, Proctor G, Spudich G, Almeida-King J, Staines D, Derwent P, Kerhornou A, Kersey P, Flicek P (2011) Ensembl BioMarts: a hub for data retrieval across taxonomic space. Database bar030.

Laemmli UK (1970) Cleavage of structural proteins during the assembly of the head of bacteriophage T4. Nature 227: 680–685.

Leonardi GA, Carlos NA, Mazzafera P, Balbuena TS (2015) *Eucalyptus urograndis* stem proteome is responsive to short-term cold stress. Genet. Mol. Biol. 38: 191–198.

Lichtenthaler HK, Schwender J, Disch A, Rohmer M (1997) Biosynthesis of isoprenoids in higher plant chloroplast proceed via a mevalonate-independent pathway. FEBS Lett. 400: 271–274.

Maier A, Zell MB, Maurino VG (2011) Malate decarboxylases: evolution and roles of NAD(P)-ME isoforms in species performing C_4_ and C_3_ photosynthesis. J. Exp. Bot. 62: 3061–3069.

Majeran W, Friso G, Ponnala L, Connolly B, Huang M, Reidel E, Zhang C, Asakura Y, Bhuiyan NH, Sun Q, Turgeon R, van Wijk KJ (2010) Structural and metabolic transitions of C_4_ leaf development and differentiation defined by microscopy and quantitative proteomics in maize. Plant Cell. 22: 3509–3542.

Mueller ND, Gerber JS, Johnston M, Ray DK, Ramankutty N, Foley JA (2012) Closing yield gaps through nutrient and water management. Nature 490: 254–257.

Pan R, Hu J (2018) Proteome of plant peroxisomes. In: Proteomics of peroxisomes. Subcellular biochemistry. del Río L, Schrader M (eds.). Springer Singapore: 3–45.

Pick TR, Bräutigam A, Schlüter U, Denton AK, Colmsee C, Scholz U, Fahnenstich H, Pieruschka R, Rascher U, Sonnewald U, Weber AP (2011) Systems analysis of a maize leaf developmental gradient redefines the current C_4_ model and provides candidates for regulation. Plant Cell. 23: 4208–4220.

Rao X, Lu N, Li G, Nakashima J, Tang Y, Dixon RA (2016) Comparative cell-specific transcriptomics reveals differentiation of C_4_ photosynthesis pathways in switchgrass and other C_4_ lineages. J. Exp. Bot. 67: 1649–1662.

Sage RF (2004) The evolution of C_4_ photosynthesis. New Phytol. 161: 341–370.

Saigo M, Alvarez CE, Andreo CS, Drincovich MF (2013a) Plastidial NADP-malic enzymes from grasses: Unraveling the way to the C_4_ specific isoforms. Plant Physiol. Biochem. 63: 39–48.

Saigo M, Bologna FP, Maurino VG, Detarsio E, Andreo CS, Drincovich MF (2004) Maize recombinant non-C_4_ NADP-malic enzyme: a novel dimeric malic enzyme with high specific activity. Plant Mol. Biol. 55: 97–107.

Saigo M, Tronconi MA, Gerrard Wheeler MC, Alvarez CE, Drincovich MF, Andreo CS (2013b) Biochemical approaches to C_4_ photosynthesis evolution studies: the case of malic enzymes decarboxylases. Photosynth. Res. 117: 177–187.

Schlüter U, Weber APM (2020) Regulation and evolution of C_4_ photosynthesis. Annu. Rev. Plant Biol. 71: 183–215.

Schulze ED, Hall AE (1982) Stomatal responses, water loss and CO_2_ assimilation rates of plants in contrasting environments. In: Physiological plant ecology II: water relations and carbon assimilation. Lange OL, Nobel PS, Osmond CB, Ziegler H (eds), Springer-Verlag: 181–230.

Schwacke R, Ponce-Soto GY, Krause K, Bolger AM, Arsova B, Hallab A, Gruden K, Stitt M, Bolger ME, Usadel B (2019) MapMan4: a refined protein classification and annotation framework applicable to multi-omics data analysis. Mol. Plant. 12: 879–892.

Shih PM, Occhialini A, Cameron JC, Andralojc PJ, Parry MAJ, Kerfeld CA (2016) Biochemical characterization of predicted Precambrian RuBisCO. Nat. Commun. 7: 10382.

Soll J, Schultz G, Rüdiger W, Benz J (1983) Hydrogenation of geranylgeraniol: two pathways exist in spinach chloroplasts. Plant Physiol. 71: 849–854.

Sweetlove LJ, Beard KFM, Nunes-Nesi A, Fernie AR, Ratcliffe RG (2010) Not just a circle: flux modes in the plant TCA cycle. Trends Plant Sci. 15: 462–470.

Titus DE, Becker WM (1985) Investigation of the glyoxysome-peroxisome transition in germinating cucumber cotyledons using double-label immunoelectron microscopy. J. Cell Biol. 101: 1288–1299.

Tronconi MA, Andreo CS, Drincovich MF (2018) Chimeric structure of plant malic enzyme family: different evolutionary scenarios for NAD- and NADP-dependent isoforms. Front. Plant Sci. 9: 565.

Tronconi MA, Fahnenstich H, Gerrard Weehler MC, Andreo CS, Flügge UI, Drincovich MF, Maurino VG (2008) Arabidopsis NAD-malic enzyme functions as a homodimer and heterodimer and has a major impact on nocturnal metabolism. Plant Physiol. 146: 1540–1552.

Tronconi MA, Gerrard Wheeler MC, Maurino VG, Drincovich MF, Andreo CS (2010a) NAD-malic enzymes of *Arabidopsis thaliana* display distinct kinetic mechanisms that support differences in physiological control.Biochem. J. 430: 295–303.

Tronconi MA, Hüdig M, Schranz ME, Maurino VG (2020) Independent recruitment of duplicated β-subunit-coding NAD-ME genes aided the evolution of C_4_ photosynthesis in Cleomaceae. Front. Plant Sci. 11: 1–15.

Tronconi MA, Maurino VG, Andreo CS, Drincovich MF (2010b) Three different and tissue-specific NAD-malic enzymes generated by alternative subunit association in *Arabidopsis thaliana*. J. Biol. Chem. 285: 11870–11879.

Tyanova S, Temu T, Sinitcyn P, Carlson A, Hein MY, Geiger T, Mann M, Cox J (2016) The Perseus computational platform for comprehensive analysis of (prote)omics data. Nat. Methods 13: 731–740.

Van Eck J (2018) The status of *Setaria viridis* transformation: *Agrobacterium*-mediated to floral dip. Front. Plant Sci. 9: 652.

Van Eck J, Swartwood K (2015) Setaria viridis. Methods Mol. Biol. 1223: 57–67.

Vranová E, Coman D, Gruissem W (2013) Network analysis of the MVA and MEP pathways for isoprenoid synthesis. Annu. Rev. Plant Biol. 64: 665–700.

Wang L, Wang C, Liu X, Cheng J, Li S, Zhu JK, Gong Z (2019) Peroxisomal β-oxidation regulates histone acetylation and DNA methylation in *Arabidopsis*. Proc. Natl. Acad. Sci. USA 116: 10576–10585.

Wang X, Gowik U, Tang H, Bowers JE, Westhoff P, Paterson AH (2009).Comparative genomic analysis of C_4_ photosynthetic pathway evolution in grasses. Genome Biol. 10: R68.

Wang Y, Bräutigam A, Weber APM, Zhu XG (2014) Three distinct biochemical subtypes of C_4_ photosynthesis? A modelling analysis. J. Exp. Bot. 65: 3567–3578.

Washburn JD, Schnable JC, Conant GC, Brutnell TP, Shao Y, Zhang Y, Ludwig M, Davidse G, Pires JC (2017) Genome-guided phylo-transcriptomic methods and the nuclear phylogentic tree of the Paniceae grasses. Sci. Rep. 7: 13528.

